# Rupture of nuclear envelope in starfish oocytes proceeds by F-actin-driven segregation of pore-dense and pore-free membranes

**DOI:** 10.1101/480434

**Authors:** Natalia Wesolowska, Pedro Machado, Ivan Avilov, Celina Geiss, Hiroshi Kondo, Masashi Mori, Yannick Schwab, Péter Lénárt

## Abstract

The nucleus of oocytes, traditionally referred to as the germinal vesicle, is unusually large and its nuclear envelope (NE) is densely packed with nuclear pore complexes (NPCs) stockpiled for embryonic development. We have shown that breakdown of this specialized NE during meiosis of starfish oocytes is mediated by an Arp2/3-nucleated F-actin ‘shell’, in contrast to microtubule-driven tearing in somatic cells. The detailed mechanism of how the cytoskeletal forces disrupt the NE remains poorly understood in any system. Here, we address the mechanism of F-actin-driven NE rupture by using live-cell and correlated super-resolution light and electron microscopy. We show that actin is nucleated within the lamina and sprouts filopodia-like spikes towards the nuclear membranes forcing lamina and nuclear membranes apart. These F-actin spikes protrude pore-free nuclear membranes, whereas the adjoining membrane stretches accumulate packed NPCs associated with the still-intact lamin network. NPC conglomerates sort into a distinct tubular-vesicular membrane network, while breaks appear in pore-free, ER-like regions. Together, our work reveals a novel function for Arp2/3-mediated membrane shaping in NE rupture that is likely to have broad relevance in regulating NE dynamics in diverse other contexts such as nuclear rupture frequently observed in cancer cells.

## Introduction

The nuclear envelope (NE), composed of inner and outer nuclear membranes, is a specialized sub-compartment of the endoplasmic reticulum (ER) that separates nucleus and cytoplasm in eukaryotic cells. The inner and outer NE is fused at nuclear pore complexes (NPCs) to mediate nucleo-cytoplasmic transport. This complex NE membrane structure is mechanically supported by a network of intermediate filaments, the lamina, lining its nucleoplasmic side (Burke and Ellenberg, 2002).

Across species and cell types there is a considerable diversity of nuclear structure in adaptation to the physiological function. For example, the composition of the lamina is adapted to provide the required high mechanical stability in muscle cells, or sufficient flexibility in immune cells, which need to squeeze through confined spaces (Thiam et al., 2016). Oocytes have a very specialized nuclear architecture with an exceptionally large nucleus, also known as germinal vesicle, storing nuclear components to support early embryonic development. The oocyte NE is thus densely packed with NPCs serving as a stockpile of these complexes (rendering oocytes a popular model to study NPCs), whereas the lamina is thick to mechanically support this very large structure (Goldberg and Allen, 1995).

The NE needs to be dismantled at the onset of every division to give microtubules access to chromosomes, and then reassembled at the end of division once the chromosomes are segregated. Dependent on the species and nuclear architecture, there is a broad diversity in disassembly mechanisms. In *Drosophila* and *C. elegans* embryos the NE and the lamina remains partially intact, whereas in vertebrates and deuterostomes (including the echinoderm starfish), the complex NE structure is fully disassembled during division. In somatic mammalian cells NE disassembly involves the complete dismantling of the NPCs, depolymerization of the lamina, and re-absorption of the nuclear membranes into the ER (Hetzer, 2010; Ungricht and Kutay, 2017).

It is common to all species in which NEBD has been investigated in detail, including somatic cells and oocytes of various species, that NEBD begins with a partial permeabilization of the NE due to phosphorylation-driven disassembly of the NPCs and other NE components (Dultz et al., 2008; Mühlhäusser and Kutay, 2007; Terasaki et al., 2001; Lénárt et al., 2003; Martino et al., 2017; Linder et al., 2017). This allows proteins up to ~70 kDa to leak in or out of the nucleus (Lénárt et al., 2003). Furthermore, it is likely that the mechanical properties of the NE are affected, i.e. the NE is weakened and destabilized as a result of phosphorylation of lamins and lamina-associated proteins (Ungricht and Kutay, 2017). Importantly however, during this first phase the overall structure of the NE, as seen by electron microscopy (EM) is still intact, and compartmentalization of large protein complexes (e.g. ribosomes, microtubules) is maintained (Terasaki et al., 2001; Lénárt et al., 2003).

It is also common to all cases that have been investigated, that the slow, phosphorylation-driven weakening of the NE is followed by a sudden rupture of the NE leading to rapid and complete mixing of cyto- and nucleoplasm. As this dramatic change is easily visible even by transmitted light microscopy, this second step is commonly identified as ‘NEBD,’ marking the transition between prophase and prometaphase of cell division. Observations from several cell types suggest that this sudden rupture requires mechanical force to be generated by the cytoskeleton. In cultured mammalian cells, microtubules tear the NE in a dynein-dependent process (Beaudouin et al., 2002; Salina et al., 2002). By contrast, we have shown recently that in the large oocyte nucleus, the actin rather than the microtubule cytoskeleton is required for NE rupture. A transient F-actin ‘shell’ is polymerized by the Arp2/3 complex on the inner surface of the NE and upon its passage, within less than two minutes, membranes undergo complete rupture (Mori et al., 2014). Critically, in neither case, i.e. in the microtubule-driven rupture in somatic cells, nor the F-actin shell in oocytes, was it understood how cytoskeletal forces mediate this rapid and dramatic reorganization of the NE.

Here, we use a combination of live-cell and super-resolution light microscopy, and correlated electron microscopy to capture these sudden changes in NE organization. We find that the F-actin shell is nucleated within the still-intact lamina and projects filopodia-like spikes into the nuclear membranes. The resulting nuclear membrane protrusions are free of NPCs, but are juxtaposed by NPC clusters. Subsequently, these NPC conglomerates invaginate and sort into NPC-rich membrane network, while breaks appear on the pore-free regions.

## Results

### Lamina remains intact during NE rupture

We have shown previously that the F-actin shell mediates NE rupture (Mori et al., 2014). However, our live-cell imaging assays lacked the resolution to visualize structural details of this dramatic reorganization occurring at the sub-micrometer scale. Therefore, we optimized staining and imaging protocols, and we developed an antibody against the only identified starfish lamin protein. This, together with the pan-NPC antibody mAb414 enabled us to co-visualize endogenous NE components together with phalloidin-stained F-actin at high resolution.

The F-actin shell is very transient, polymerizing and depolymerizing within 2 minutes that also necessitated development of a reliable temporal reference for fixed-cell assays. Fortunately, the F-actin shell emerges in a highly reproducible spatial pattern, which enabled us to time the fixed samples by correlating them with morphologies observed live (compare Fig. 1A and B). The F-actin shell first appears on the inner side of NE as an equatorial band of foci when the NE is still intact and impermeable to large dextrans (Fig. 1A, 0 s). Then, as these F-actin foci grow and intensify, merging into a continuous F-actin shell, the first breaks on the NE appear, visualized by localized entry of large dextrans (Fig. 1A, 45 s). The shell then spreads towards the poles, followed by a wave of membrane rupture with a ~30 s delay (Mori et al., 2014). Using these distinct morphological features, we were able to reliably identify and order stages of NE rupture in fixed samples.

**Figure 1.**
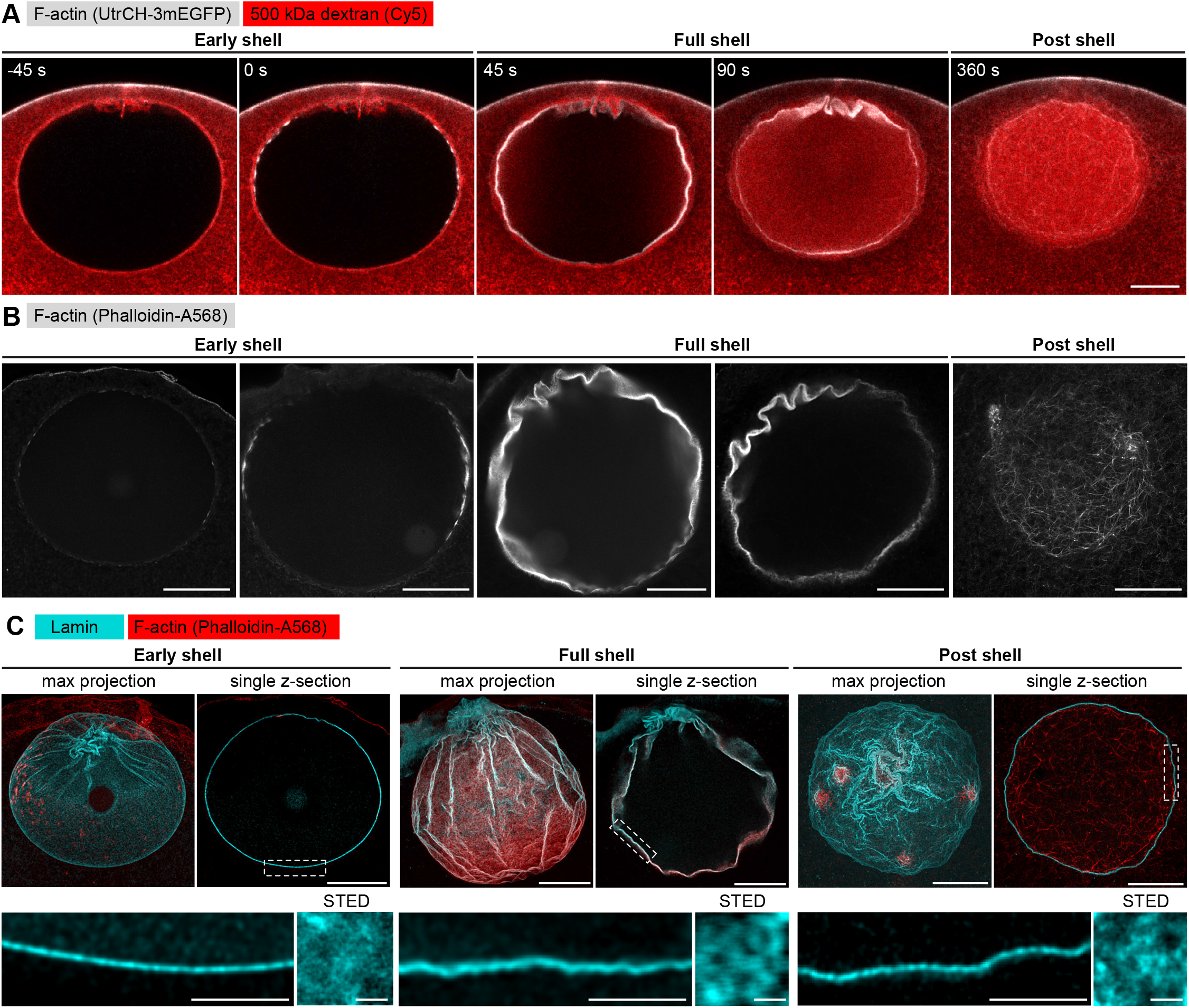
The lamina remains intact during NE rupture. **(A)** Live imaging of UtrCH-3mEGFP (white) and Dextran-500 kDa (red) in a starfish oocyte undergoing NEBD. Selected single confocal sections are shown from a time series; scale bar is 20 μm. **(B)** Fixed samples with F-actin labeled by phalloidin-AlexaFluor568. Individual confocal sections are shown and ordered to match the live time series in (A). Scale bar is 20 μm. **(C)** Immunostained starfish oocytes with anti-lamin antibody shown in cyan and phalloidin-AlexaFluor568 in red. Shown are three time-points: early shell, full shell and post shell. Each panel shows a maximum projection of the whole z-stack (left), a single selected optical section at the equatorial plane (right), and a zoom of the area in the single section highlighted with dashed rectangle (bottom). The small insets on the bottom right show *en face* views of the lamina in oocytes stained with the anti-lamin antibody and imaged by STED at the corresponding stages. Scale bars are 20, 5 and 0.5 μm, respectively.

With this assay in hand, we first wanted to clarify whether the F-actin shell destabilizes the NE by tearing the lamin network, as it has been reported in somatic cells for microtubule-driven NE rupture (Beaudouin et al., 2002; Salina et al., 2002). We have addressed this question earlier in starfish oocytes, however, at that time we had exogenously overexpressed a GFP fusion of human lamin B, which could have had different disassembly kinetics (Lénárt et al., 2003). To directly visualize the endogenous lamina this time we analyzed samples stained with the starfish anti-lamin antibody at different stages of NE rupture. This confirmed our previous observations that even minutes after NE rupture the endogenous lamin network remains continuous, although it folds and ruffles as the nucleus collapses during NEBD (Fig. 1C). Additionally, imaging portions of the lamina *en face* by stimulated emission depletion (STED) microscopy suggests that the lamin mesh gradually coarsens during the process of NEBD (Fig. 1C).

We conclude that the rupture of the NE does not proceed by F-actin-induced tearing or rapid disassembly of the lamina, which remains a continuous network throughout NEBD.

### The F-actin shell assembles within the lamina sprouting spikes that detach nuclear membranes

In order to localize the F-actin shell relative to NE components, we next co-localized the lamina or the nuclear membranes (as marked by NPCs) at the time of shell formation with the F-actin shell stained by phalloidin. We observed that while the lamina co-localized with phalloidin, the NPC staining formed a separate layer of fragmented appearance up to 500 nm ‘above’ the F-actin shell (Fig. 2A, B). Thus, the still-intact lamina appears to serve as the scaffold upon which the F-actin shell assembles, whereas the nuclear membranes appear to fragment and detach from the lamina.

**Figure 2.**
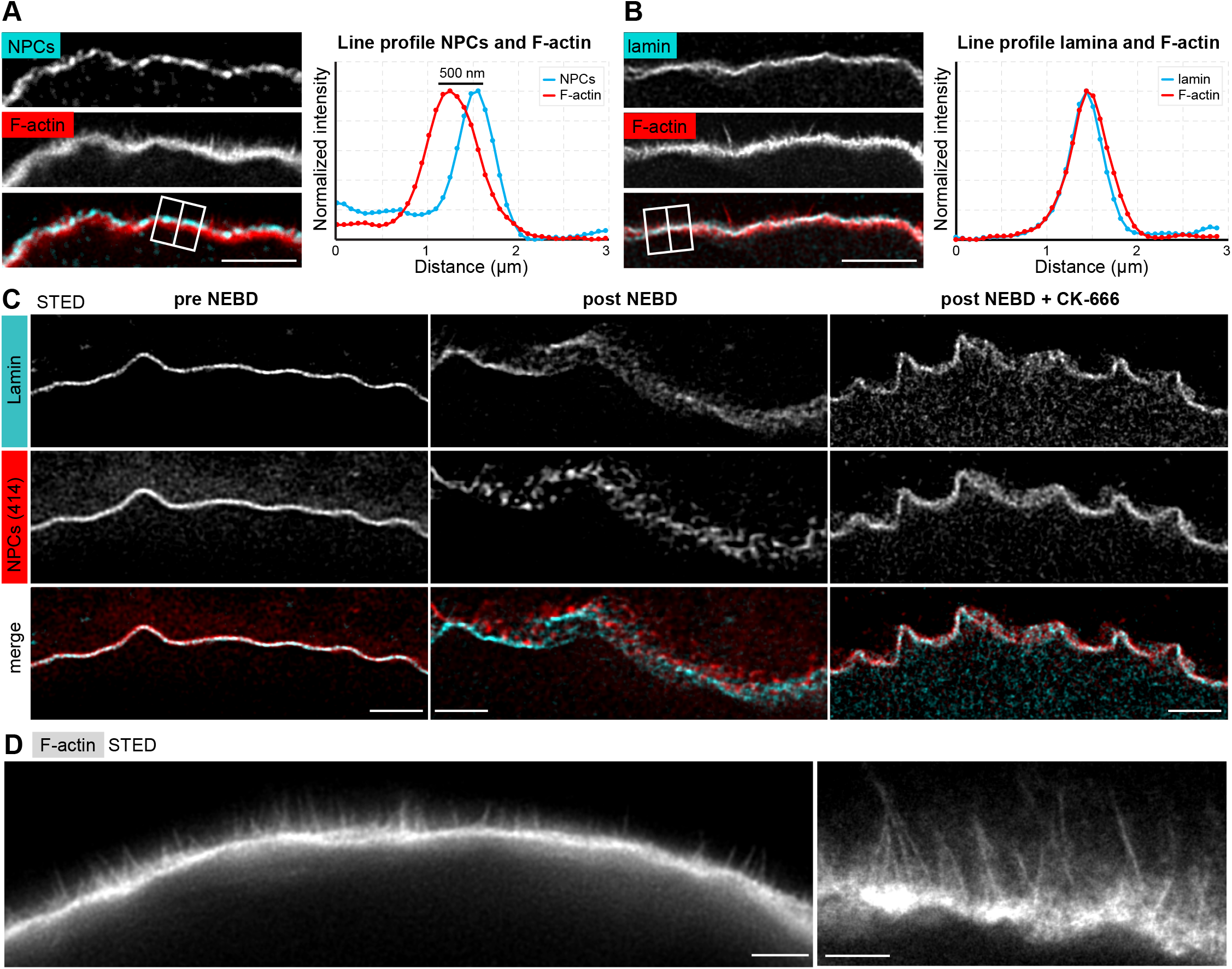
The F-actin shell separates lamina and nuclear membranes during NEBD. **(A)** Left: Portion of the NE undergoing rupture immunostained with mAb414 for NPCs (cyan) and phalloidin-AlexaFluor568 for F-actin (red). Shown is a crop of a portion of NE from a confocal Z-section; scale bar 2 μm. Right: Plot of a line profile over the region marked with a white rectangle; normalized intensities of both channels are shown. **(B)** Same as (A), except stained with anti-lamin antibody and phalloidin-AlexaFluor568. **(C)** Portions of the NE stained with anti-lamin antibody (cyan) and mAb414 (red) and imaged by STED. Left: before NEBD, middle: after NEBD, right: after NEBD but first treated with CK-666 to inhibit the formation of the F-actin shell. Scale bars are 2 μm. **(D)** Phalloidin-Alexa568 staining of F-actin shell imaged by STED microscopy. Scale bars are 5 and 2 μm, respectively.

We confirmed these observations by STED imaging of the lamina and NPCs (note that these samples were co-stained with phalloidin for staging, but it was not possible to image the third color on this particular STED setup). Before NE rupture, the NPC and lamina stainings overlapped at the approx. 50 nm resolution afforded by STED, what is consistent with the known ultrastructure of the NE (Fig. 2C, pre-NEBD) (Burke and Ellenberg, 2002). In stark contrast, at the shell stage we could visualize NPC-stained NE fragments “floating” above the intact lamina (Fig 2C, post-NEBD). This detachment is dependent on the F-actin shell, because when we prevented F-actin shell formation by inhibiting Arp2/3 by the small molecule inhibitor, CK-666 lamina and NPCs did not separate (Fig. 2C, post-NEBD + CK-666). In these images the NE appears ruffled due to collapse of nuclear volume, but remains intact, consistent with earlier findings (Mori et al., 2014).

Additionally, STED imaging of samples optimally fixed for phalloidin staining, revealed filopodia-like F-actin spikes of 0.5-2 μm in length, spaced ~0.1 μm, and extending from the base of the F-actin shell towards nuclear membranes (Fig 2D).

Taken together, our data show that the F-actin shell nucleates in the lamina and extends filopodia-like spikes towards the nuclear membranes. In this process the nuclear membranes fragment and separate from the lamina. However, a critical issue in interpreting these results is that we rely on the NPC staining to represent nuclear membranes. As the oocyte NE in interphase is fully packed with an almost crystalline array of NPCs (Lénárt et al., 2003), this assumption is reasonable, but during NE rupture exactly this arrangement may change. Unfortunately, our efforts to visualize membranes by light microscopy remained futile, because preserving F-actin in fixed oocytes requires addition of detergents to the fixative, which is incompatible with preserving fine membranous structures. Thus, we turned to correlated electron microscopy.

### Correlative EM captures intermediates of NE rupture

In order to clarify the F-actin mediated rearrangements of nuclear membranes, we decided to target early stages of the F-actin shell formation, i.e. the approx. 30 s time-window, when parts of the NE are already ruptured, while other regions are still intact. We expected that at this stage we could observe intermediate steps of NE rupture, spatially arranged within the same sample.

For this purpose we developed a correlative electron microscopy protocol using high-pressure freezing and freeze substitution, resulting in an excellent preservation of cellular structures (Burdyniuk et al., 2018). Correlation to light microscopy was achieved by using fluorescently-labeled dextrans, which are directly visible on EM sections, as indicators of NEBD progression (Fig. 3B, C): the small, 25 kDa dextran enters the nucleus already in the first phase of NEBD through the disassembling NPCs, while the large, 160 kDa dextran only enters when the NE is ruptured (Lénárt et al., 2003). Thus, when the 25 kDa dextran almost completely fills the nucleus, but the large, 160 kDa dextran is still excluded identifies the time-window of F-actin shell formation and NE rupture.

**Figure 3.**
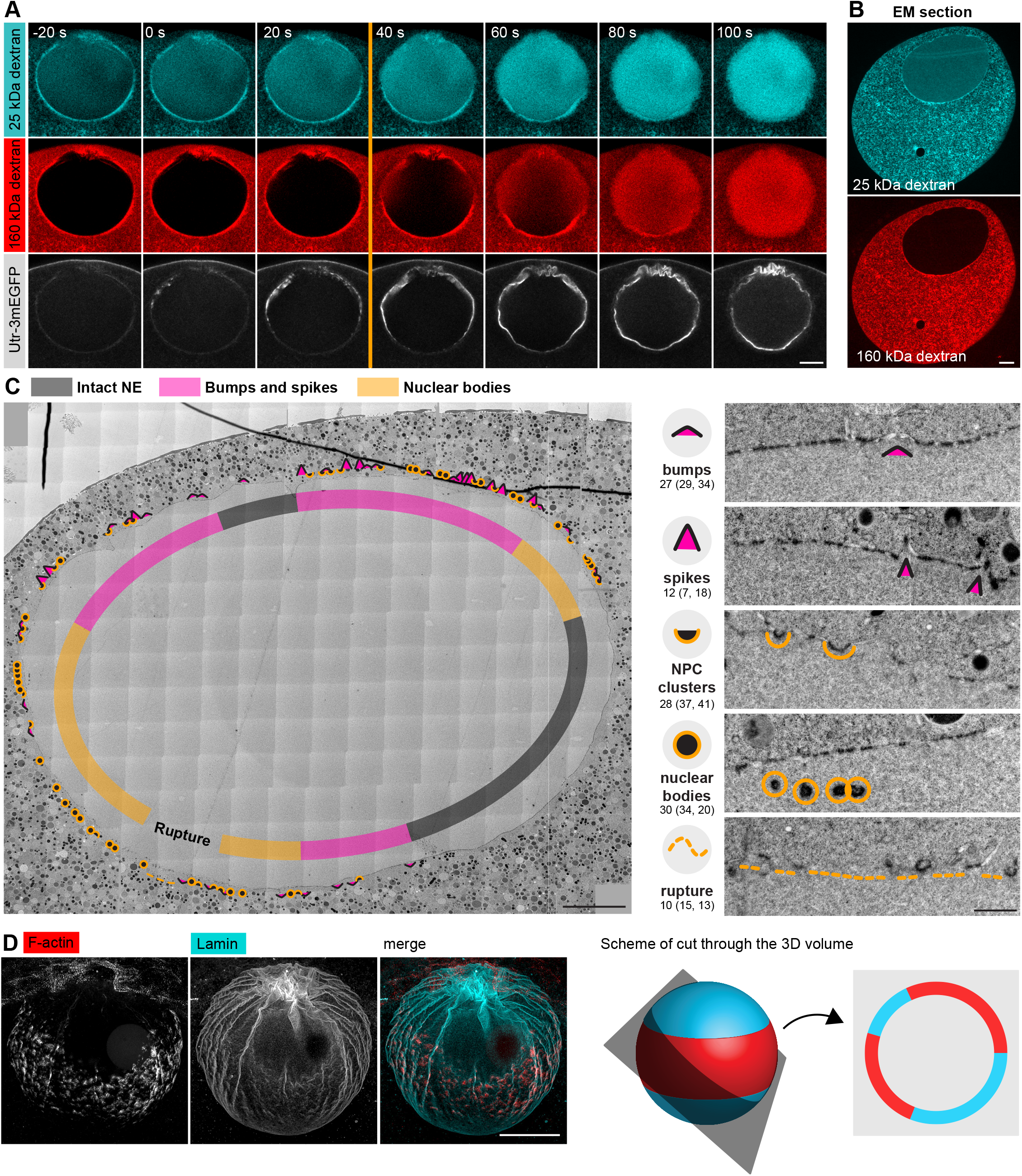
Correlative EM approach captures NEBD intermediates. **(A)** Live imaging of a starfish oocyte undergoing NEBD and injected with a 25 kDa Cy5-labeled dextran (cyan) 160 kDa TRITC-labeled dextran (red) and UtrCh-3mEGFP (white). Shown are selected 2-frame projections from a confocal time-series imaged at 1 frame per second. Orange line marks the moment right prior to rupture of the membrane, corresponding to the predicted time for the EM sample shown in (B) and (C). Scale bar is 20 μm. **(B)** Wide-field fluorescence image of a 70 nm section of a Lowicryl-embedded oocyte undergoing NEBD and injected with 25 kDa Cy5-labeled dextran (cyan) and 160 kDa TRITC-labeled (red) dextran. Scale bar is 20 μm. **(C)** A whole-nucleus tile of transmission EM images stitched automatically for a section from oocyte shown in B. Symbols around the nucleus correspond to NE rupture intermediates observed. Symbol legend with examples (crops from the tiled image) is shown to the right. Under each symbol, numbers correspond to the count of these events in the section shown, and in parentheses the count in two adjacent sections (shown in Fig. S1-3). The band tracing the NE within the nuclear space demarcates areas with color-code for predominant membrane features. For full resolution image, see Fig. S1. Scale bar is 10 μm. **(D)** An oocyte fixed and stained with anti-lamin antibody (cyan) and phalloidin-AlexaFluor568 at early shell formation. Shown is a maximal Z projection. Right: a scheme illustrating the 3D geometry of the EM section. Scale bar: 20 μm.

We then performed automated large field-of-view montage transmission EM imaging of the whole nuclear cross-section to assess the state of the NE. An overview is shown in Fig. 3C, this and two additional montages are available at high resolution as Supplemental Data (Fig. S1-3). The montage illustrates the key advantage of the system, whereby the progression of NE rupture can be observed spatially ordered on a single section of the large oocyte nucleus. The arrangement of the rupture site is fully consistent with live and fixed light microscopy data: NE rupture initiates near the ‘equator’ of the nucleus spreading as a wave towards the poles (Fig. 3D).

We carefully examined these large montages and observed a set of frequently recurring characteristic membrane configurations. We defined them into four categories and assigned each a symbol to mark their incidence (Fig. 3C). Numbers under each category quantify the occurrence of each feature within the section, and numbers in parentheses represent similar quantification in two other sections shown in the Supplement, illustrating that these are frequent structures at the time of NE rupture.

Taken together, using our correlative light and electron microscopy approach, we were able to capture oocytes in the process of NE rupture.

### F-actin spikes protrude pore-free nuclear membranes

The large dataset, the high frequency of events observed and, importantly, the spatial arrangement from the equatorial rupture site towards the still-intact poles, allowed us to reconstruct the steps of NE rupture.

First, as consistent with earlier observations, in immature oocytes, as well as in oocytes just before NE rupture, and even in the intact polar regions of the NE undergoing rupture, the NE is smooth, continuous and is tightly packed with NPCs with a regular spacing of ~200 nm (Fig. 4A, B and S4) (Lénárt et al., 2003). In contrast, in areas closer to the rupture site we observed regions with gaps in NPC occupancy, the number and size of which increased proximal to this region (Fig. 4C). In the vicinity of the site, gaps appeared to evolve into ‘bumps’ and membrane spikes (Fig. 4D). Although reconstructing the full 3D architecture of these spikes is challenging even in 300-nm-thick tomographic sections, the most prominent spikes we observed rise ~1 μm above the level of the NE, exactly matching in size to F-actin spikes seen by phalloidin staining (Fig. 4D). These spikes contain fibrous densities fully consistent with actin filaments (Inset 3, Fig. 4D), clearly distinguishable on tomographic reconstructions (Fig. 4E). Importantly, both on thin sections and tomographic reconstructions we observed these spikes to be covered by continuous nuclear membranes almost completely free of NPCs (Fig. 4D).

**Figure 4.**
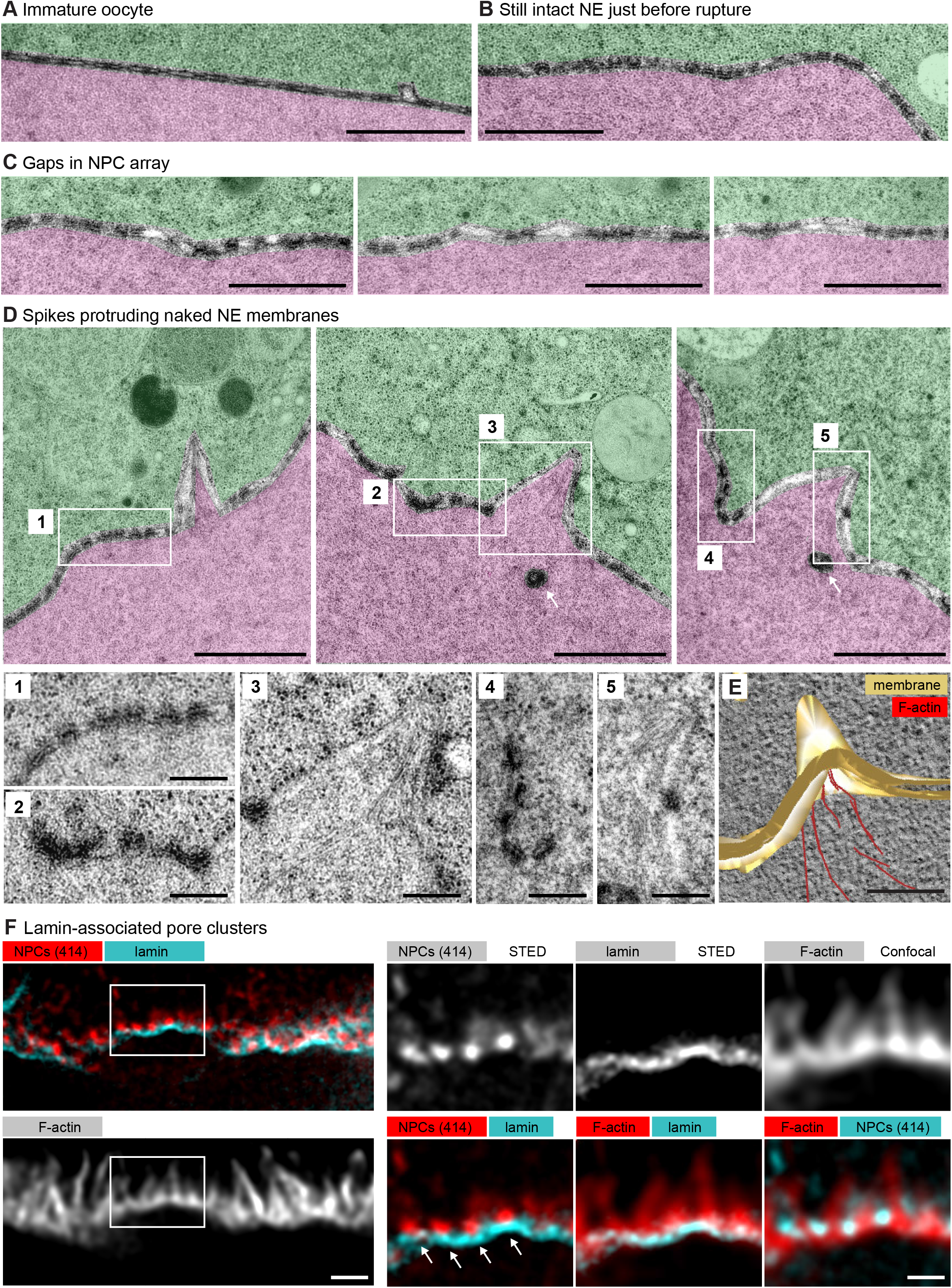
Spikes protrude bare nuclear membranes. **(A-D)** Transmission EM images from the oocyte shown in Fig. 3D showing intermediates of NE rupture. Transparency coloring distinguishes the cytoplasm (green) from nuclear area (pink) based on the presence of ribosomes. Scale bars are 1 μm, except in the zooms 1-5, where they are 250 nm. On D arrows point to nuclear bodies. Zooms of areas outlined with white rectangles are shown below. **(E)** Tomogram of a NE spike from a 300-nm thick section of the oocyte shown in Fig. 3D. with model overlay segmented manually. Scale bar is 200 nm. **(F)** STED image of the NE at the shell stage stained for NPCs (mAb414), lamina and phalloidin-AlexaFluor568. Separate channels and overlays are shown in the combinations indicated. Arrows point at nucleoplasmic bodies. Scale bar is 1 and 0.5 μm, respectively.

These pore-free areas are surrounded by NPC-dense adjacent regions with substantially reduced pore-to-pore distance of ~100 nm (Fig. 4D and S4). We are unable to estimate the 3D membrane areas from 2D sections, nevertheless our data suggest that NPC-free areas pushed out by the F-actin shell may cause the crowding of NPCs in juxtaposed regions.

These intriguing membrane morphologies prompted us to re-examine samples at the shell stage by immunofluorescence. For this purpose we further optimized our fixation and STED imaging protocols. These high resolution light microscopy data are fully consistent with observations made by EM (Fig. 4F): F-actin spikes extend out of the lamina with no NPC staining covering them except for dense NPC clusters at their base. Additionally, the light microscopy data reveals that the lamina runs at the base of the F-actin shell, and is below but apparently is still attached to NPC clusters.

Together these data suggest that the growing gaps between NPCs, which then develop into bumps and spikes, are protrusions generated by Arp2/3-driven actin polymerization pushing membranes and lamina apart. These protrusions are free of NPCs most likely because NPCs are still attached to the lamina, and thus NPCs are held back and cluster at the base of spikes.

### NPC-rich membranes sort into a tubular-vesicular network while pore-free regions rupture

Accompanying the spikes and NPC-rich clusters, we observed additional frequent membrane structures, which we call nucleoplasmic bodies: dense, round structures 200-500 nm in diameter beneath the NE in the nucleoplasm (Fig. 4D, arrows). Careful examination of intermediates suggest that nucleoplasmic bodies may form from NPC-rich clusters, initially inducing pits curving into the nucleoplasm (Fig. 5A and S4). This strongly suggests that nucleoplasmic bodies are inverted NE tubules filled with cytoplasm inside. Tomograms confirm that the electron density along the boundary of the bodies corresponds to closely juxtaposed NPCs with an intact central ring structure (Fig. 5B).

**Figure 5.**
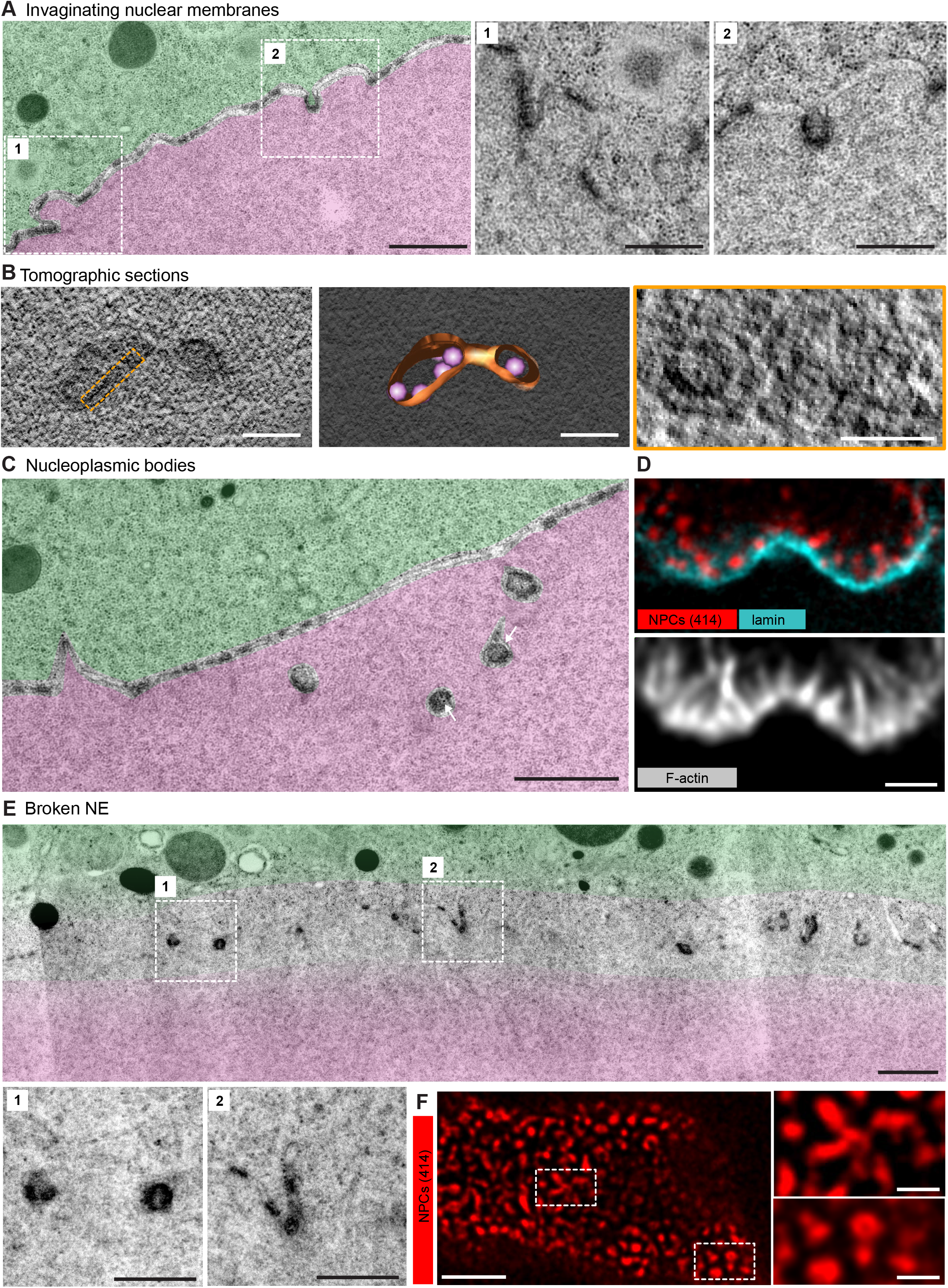
NPC clusters segregate while rupture occurs at pore-free regions. **(A)** Transmission EM images from the oocyte shown in Fig. 3D and colored as in Fig. 4A-D, showing NPC clusters membrane invaginations. Zooms of portions outlined with white rectangles are shown below without color transparencies. Scale bars are 1 μm and 500 nm, respectively. See Fig. S4 for more examples. **(B)** Tomogram of a nucleoplasmic body (left), with a model overlaid (middle). Right: re-slicing of the volume perpendicular to the view on the left corresponding to the area outlined with an orange dashed rectangle. Scale bars are 200 and 100 nm, respectively. **(C)** Transmission EM images as in (A) showing an area with nucleoplasmic bodies. Arrows point at ribosomes present within nucleoplasmic bodies. Scale bar is 1 μm. **(D)** STED image of the NE at the shell stage stained for NPCs (mAb414) (red), lamina (cyan) and phalloidin-AlexaFluor568 (gray). Scale bar is 1 μm. **(E)** Transmission EM images as in (A) showing an example of NE rupture. Zooms of the areas outlined with white squares are shown below. Scale bar is 1 μm, and 500 nm for zooms. **(F)** *En face* STED image of the NE at the shell stage stained for NPCs (mAb414) (red). Scale bar is 1 and 0.5 μm, respectively.

These bodies often appear in a beads-on-a-string arrangement with a slightly electron-denser nucleoplasmic material connecting them (Fig. 5C). Above them the NE appears to be still intact, consisting mostly of NPC-free nuclear membranes and still maintaining the nucleo-cytoplasmic boundary as judged by the distribution of ribosomes (Fig. 5C). Light microscopy of the same stages suggests that these structures may actually be related to spikes in a different geometrical arrangement: here F-actin spikes do not protrude the membrane into the cytoplasm, but rather as they grow into the membrane the nucleoplasmic bodies are pulled by the base of the spike into the nucleoplasm. Light microscopy clearly shows that the lamina runs underneath and appears to be attached to the nucleoplasmic bodies (Fig. 5D).

Finally, in areas which nucleoplasmic bodies are most frequent and exaggerated, we observe membrane rupture evidenced by presence of ribosomes in the nuclear regions (Fig. 5E). In these areas nucleoplasmic bodies are neighbored by bits of NPC-packed membranes. Light microscopy of these stages reveals a complex tubular-vesicular network densely labeled by NPC staining, in which we were able to occasionally resolve cross-sections of tubules consistent with our EM data.

Together, our data evidences that NE rupture proceeds by F-actin-driven sorting of NE membranes into pore-dense and pore-free, ER-like membrane networks.

## Discussion

Here, we resolved structural intermediates of rapid NE rearrangements mediated by the transient F-actin shell by using super-resolution light microscopy and correlative EM in starfish oocytes. Based on these data we propose the following model for NE rupture (Fig. 6): The first step is formation of F-actin foci within the lamina. We hypothesize that these foci form at the time when cytoplasmic components, such as the Arp2/3 complex and actin monomers, reach a critical concentration in the nucleus as a result of the gradual, phosphorylation-driven disassembly and increasing leakiness of the NPCs in the first phase of NEBD. Once triggered, F-actin foci grow rapidly, which is expected based on the autocatalytic nature of Arp2/3-mediated nucleation of branched F-actin networks. As they spread, they merge to a continuous shell. The filaments seem to preferentially grow from the shell base in the lamina towards nuclear membranes, and push against them. This asymmetry may be explained by fact that force imposed on actin filaments promotes nucleation of branched meshwork (Bieling et al., 2016). Intriguingly, F-actin networks nucleated *in vitro* on micropatterned activated Arp2/3 show a strikingly similar morphology to the F-actin shell with filopodia-like bundles pointing away from a base of dense branched network (Reymann et al., 2010). This suggests that localized activation of Arp2/3 within the lamina may be sufficient to explain the morphology of the F-actin shell.

**Figure 6.**
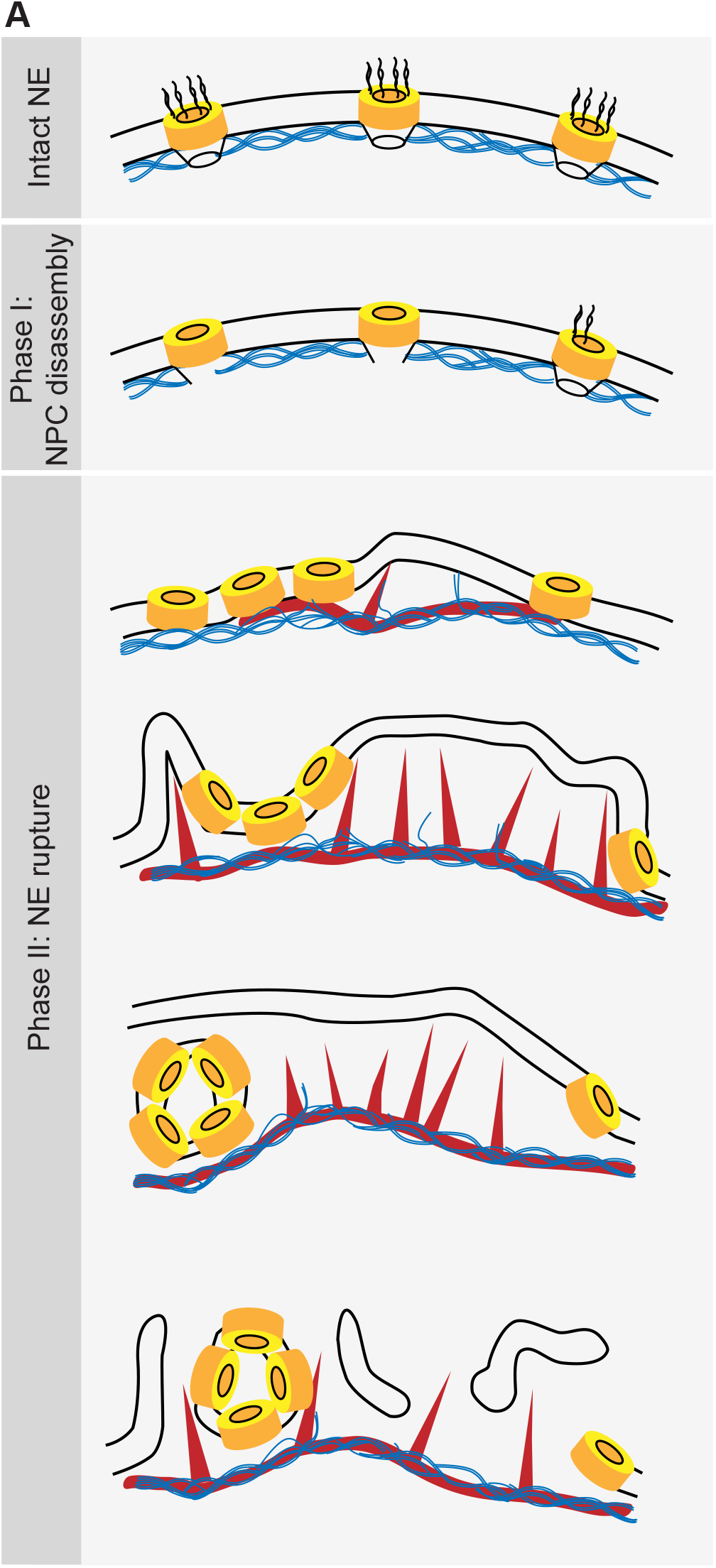
Model of F-actin-driven NE rupture. Intact NE: interphase organization of nuclear membranes (black lines) with regularly spaced NPCs (yellow cylinders) featuring cytoplasmic filaments and nuclear baskets. Nuclear baskets are embedded in the lamin network (blue filaments). Phase I of NEBD: peripheral NPC components are gradually released, but the NPC core and overall NE structure remains intact. Phase II: NE rupture. First, small patches of F-actin (red) form within the lamina. F-actin patches grow and merge to a shell pushing apart NPCs still partially anchored in the lamina. As frequent F-actin spikes further sever the lamin-to-NE attachments, NPCs segregate into conglomerates leaving stretches of unstable bare membrane, where breaks appear.

Our EM data clearly show that the F-actin shell protrudes pore-free nuclear membranes, separating these from the lamina. We propose that these membranes are cleared of NPCs, because NPCs are attached to the still-intact lamina at this stage, and thus are held back, while membranes are protruded by actin polymerization. Then, as the NPCs accumulate between pore-free spikes, membranes buckle into the nucleoplasm and invaginate to form nucleoplasmic bodies. Our data suggest that pore-free nuclear membranes separated from the lamina are unstable, and thus rupture and rearrange into an ER-like reticular structure. Although this hypothesis remains rather speculative at this point, it is intriguing to consider that repair of interphase NE rupture, as well as reassembly of NE after division, involves the ESCRT machinery (Denais et al., 2016; Raab et al., 2016; Olmos et al., 2015). In our NEBD intermediates we observe similar membrane topologies to those involved in NE repair, suggesting that the ESCRT complex may also be involved in these membrane rearrangements.

In agreement with our model, it has been shown in various other physiological contexts that detaching the nuclear membranes from the lamin network leads to NE rupture. For example, NE rupture frequently occurs in cancer cells, in particular in micronuclei, where breaching of the NE barrier is preceded by local lamin disruption (Hatch et al., 2013). More generally, recent work shows that the “breakability” of the nucleus in migrating cells is dependent on lamin composition (Thiam et al., 2016; Davidson and Lammerding, 2014; Denais et al., 2016). Indeed, in somatic cells it has been show that NEBD proceeds by microtubule-mediate tearing of the lamina (Beaudouin et al., 2002; Salina et al., 2002), but the precise morphology of the NE intermediates at a resolution comparable to what we could achieve in starfish oocytes is not known. Therefore, it is well possible that a similar mechanism based on segregation of lamina, NPCs and membranes plays a critical role in destabilizing the NE during NE rupture in somatic cells as well.

As mentioned in the Introduction, there is wide diversity across species and cell types in the extent of NE disassembly during division. For example, the lamina persists much longer in many species as compared to mammalian somatic cells (Gruenbaum et al., 2003). Intriguingly, an F-actin shell similar to that in starfish oocytes has been observed in several species, although these structures are yet to be characterized in detail. This includes other echinoderm species such as sea urchin (Burkel et al., 2007), the cnidarian model *Nematostella vectensis* (DuBuc et al., 2014), as well as polychaete worms (Jacobsohn, 1999). These examples strongly suggest that F-actin mediated NE rupture may be widely spread across animal species, and we speculate that the presence or absence of this mechanism may be correlated with differences in NE disassembly dynamics, nuclear size and other differences in physiology.

Finally, the clearing of NPCs off the membrane appears to be conserved to NEBD in organisms with partially open mitosis. Tearing of the NE has been observed in the fungus *Ustilago maydis* and the yeast *Schizosaccharomyces japonicus*. In *U. maydis* the shearing is caused by microtubules pulling the nucleus through a small opening into the bud of the daughter cell (Straube et al., 2005). *Sz. japonicus,* on the other hand, splits the NE by stretching the nucleus between the two poles of a dividing cell (Aoki et al., 2011). Thus, NE rupture occurs by different means, but intriguingly both show evidence of clearing NPCs before the NE is torn: *Sz. japonicus* redistributes the NPCs to the two poles freeing naked membranes at the NE regions destined to be broken. NPCs of *U. maydis* on the other hand initiate release of nucleoporins prior to rupture just as in higher eurkaryotes (Straube et al., 2005; Aoki et al., 2011; Theisen et al., 2008).

Taken together, our data suggest that the F-actin shell destabilizes the NE by segregating pore-dense and pore-free membranes, providing the first mechanistic explanation for the sudden collapse of the NE structure during its breakdown. As discussed above, this mechanism is likely to function is many animal species, and while in other species forces may be generated by other means than Arp2/3-mediated actin polymerization, segregating nuclear membranes from the lamin network appears to be a general feature of nuclear rupture observed in dividing mammalian somatic cells, as well as during interphase NE rupture frequent in cancer cells.

## Materials and methods

### Oocyte collection and injection

Starfish (*Patiria miniata*) were obtained in the springtime from Southern California (South Coast Bio-Marine LLC, Monterey Abalone Company or Marinus Scientific Inc.) and kept at 16°C for the rest of the year in seawater aquariums of EMBL’s marine facilities. Oocytes were extracted from the animals fresh for each experiment as described earlier (Lénárt et al., 2003). mRNAs and other fluorescent markers were injected using microneedles, as described previously (Jaffe and Terasaki, 2004; Borrego-Pinto et al., 2016). mRNA was injected the day before to allow protein expression, while fluorescently labeled protein markers or dextrans were injected a few hours prior to imaging. Meiosis was induced at initiation of experiment by addition of 1-methyladenine (1-MA, 10 μM, Acros Organics). NEBD normally started at 20-25 minutes after 1-MA addition.

### Fluorescent markers and antibodies

To label F-actin, 3mEGFP-UtrCH mRNA was synthesized *in vitro* from linearized DNA templates using the AmpliCap-Max T7 High Yield Message Maker kit (Cellscript), followed by polyA-tail elongation (A-Plus Poly(A) Polymerase Tailing Kit, Cellscript). mRNAs were dissolved in water (typical concentration 3-5 μg/μl) and injected into the oocyte up to 5% of the oocyte volume.

Phalloidin labeled with the indicated Alexa fluorophores (Invitrogen) was dissolved in methanol, and was then air-dried prior to use and dissolved in PBS for immunostaining.

For dextrans, amino-dextrans were labeled with succinimidyl ester dye derivatives (Cy5) or purchased already in labeled form (TRITC), purified and injected into oocytes as described earlier (Lénárt et al., 2003).

The pan-NPC antibody mAb414 was purchased from BioLegend or Sigma (catalogue #902907, #N8786, respectively). To produce the anti-starfish-lamin antibody, first the *Patiria miniata* lamin sequence was identified by BLAST searches in our transcriptome-based database (http://www.lenartlab.embl.de:4567/) by comparisons to the human lamin B amino acid sequence, and confirmed by reverse searches to other species. Furthermore, the corresponding mRNA was expressed as a mEGFP fusion and showed the expected localization to the NE in starfish oocytes (not shown). Peptide antibodies were then produced against the “histone-interaction peptide” region of starfish lamin (GTKRRRLDEEESMVQSS), which was used as the antigen for rabbit immunization. Antibody production and affinity purification was performed by Cambridge Research Biochemicals. The antibody’s specificity was confirmed in Western blot showing an expected-sized band and immunostaining showing localization to the nuclear rim in starfish oocytes.

### Immunostaining

Oocytes were fixed at desired times in a PFA/GA fixative (100 mM HEPES pH 7.0, 50 mM EGTA, 10 mM MgSO_4_, 0.5% Triton-X100, 1 or 2% formaldehyde, 0.2 or 0.4% glutaraldehyde) modified from Strickland et al. (Strickland et al., 2004). Active aldehyde groups remaining post fixation were quenched by 0.1% solution of NaBH_4_ or 200 mM NH_4_Cl and 200 mM glycine. Subsequently, samples were permeabilized and blocked in PBS+0.1% Triton-X100 plus 3% BSA and the Image-IT reagent (ThermoFisher Scientific). Antibody staining was done overnight for the primary antibody and for 2-3 h for the secondary antibody in PBS+0.1% Triton-X100 at room temperature. Oocytes were mounted with the antifade agent ProLongGold (ThermoFisher Scientific) under a coverslip pressed quite firmly onto tiny pillars of grease or double sided tape (Scotch).

### Light microscopy

Live cell movies were acquired on a Leica SP5 confocal microscope using a 40x HCX PL AP 1.10 NA water immersion objective lens (Leica Microsystems). Fixed oocytes were imaged on a Leica SP8 microscope equipped with the HC PL APO 1.40 NA 100x oil immersion objective according to Nyquist criteria. For STED imaging, suitable Abberior STAR 580 and Abberior STAR RED or Abberior STAR 635P secondary antibodies or nanobodies were used (Abberior, NanoTag). Samples were imaged on a Leica SP8 STED microscope, with the HC PL APO CS2 1.40 NA 100x oil immersion objective and using the 775 nm depletion laser. Alternatively we used an Abberior Instruments STEDYCON scan head mounted onto a Nikon Ti2 microscope equipped with 100x CFI Plan Apochromat Lambda NA 1.45 oil immersion objective lens, or an Abberior Instruments Expert Line STED microscope using an Olympus 100x UPLSAPO 100XS NA 1.4 oil immersion objective.

Images were deconvolved using the Huygens software (Scientific Volume Imaging) with either confocal or STED settings as appropriate.

### Electron microscopy

The protocol is described in detail in Burdyniuk et al. (Burdyniuk et al., 2018). In brief, oocytes were injected with a mixture of dextrans and a small batch was tested for meiosis timing. At the approximate time of NEBD they were transferred into a carrier (3 oocytes in 0.3 μl of sea water) and most of the water was removed with filter paper. Oocytes were immediately covered with a drop of 1-hexedecene, and immediately high-pressure frozen. Oocytes were freeze-substituted into Lowicryl HM-20. To stage the oocytes, light microscopy of EM sections was used to determine the progress of dextran entry. Selected sections were then post-stained with lead citrate and imaged using a BioTwin CM120 Philips transmission electron microscope at 120 kV. Large TEM montages were acquired using a JEOL JEM-2100Plus transmission electron microscope at 120 kV. Tomograms were reconstructed from tilt series acquired on FEI Tecnai F30 transmission electron microscope at 300 kV with 1.554 nm pixel size.

## Acknowledgements

We thank the members of the Lénárt laboratory for reagents and support, in particular Kálmán Somogyi, Andrea Callegari, Johanna Bischof, Joana Borrego-Pinto and Philippe Bun. We also thank EMBL’s Advanced Light Microscopy Facility for essential support, specifically Marko Lampe for the help with STED imaging. We thank the Electron Microscopy Core Facility, and Paolo Ronchi for sharing expertise during development of the EM protocol. We thank EMBL’s Laboratory Animal Resources and Kresimir Crnokic in particular.

Research in P.L.’s laboratory was funded by the European Molecular Biology Laboratory (EMBL) and the Deutsche Forschungsgemeinschaft (DFG) through the grant GZ LE 2926/1-1 AOBJ 603520 in frames of the Priority Programme SPP 1464. The laboratory is currently funded by the Max Planck Society.

## Author contributions

N.W. and P.L. conceived the project and designed the experiments. N.W. performed most experiments, I.A., H.K. and M.M. provided additional experimental data, and C.G. analyzed tomograms. N.W. and P.M. carried out electron microscopy under supervision of Y.S. N.W. and P.L. wrote the manuscript.

## Supplemental Material

### Supplemental Figure Legends

**Supplemental Figure 1. High-resolution TEM montage of a section through the nuclear region of oocyte undergoing NEBD (shown on Fig. 3C). Scale bar 10 μm.**

**Supplemental Figure 2. High-resolution TEM montage of a section through the nuclear region of oocyte undergoing NEBD, section adjacent to the one shown in Supplemental Figure 1. Scale bar 10 μm.**

**Supplemental Figure 3. High-resolution TEM montage of a section through the nuclear region of oocyte undergoing NEBD, section from the same oocyte as shown in Supplemental Figure 1. Scale bar 10 μm.**

**Supplemental Figure 4. Examples of NPC-rich clusters.** Shown are crops from sections shown in Figure S2 **(A)** and S3 **(B)**. The crops each show a membrane feature identified NPC clusters: a concave NE-membrane portion rich in NPCs. To help orientation, the first crop in each row was pseudo-colored for cytoplasm (green), nucleus (pink) and NPC-rich cluster (orange). Scale bar is 500 nm. **(C)** Box plot of distances between neighbor NPCs in different regions and samples. Interphase is an immature oocyte sample; pre-NEBD is a sample fixed a few minutes prior to NE rupture; the other groups come from different regions of the sample shown in Fig. 3C. Measurements were done manually by measuring the length of the line between two NPCs.

