## Supplementary figures and images for "Rupture of nuclear envelope in starfish oocytes proceeds by F-actin-driven segregation of pore-dense and pore-free membranes"

### Supplemental Figure 1

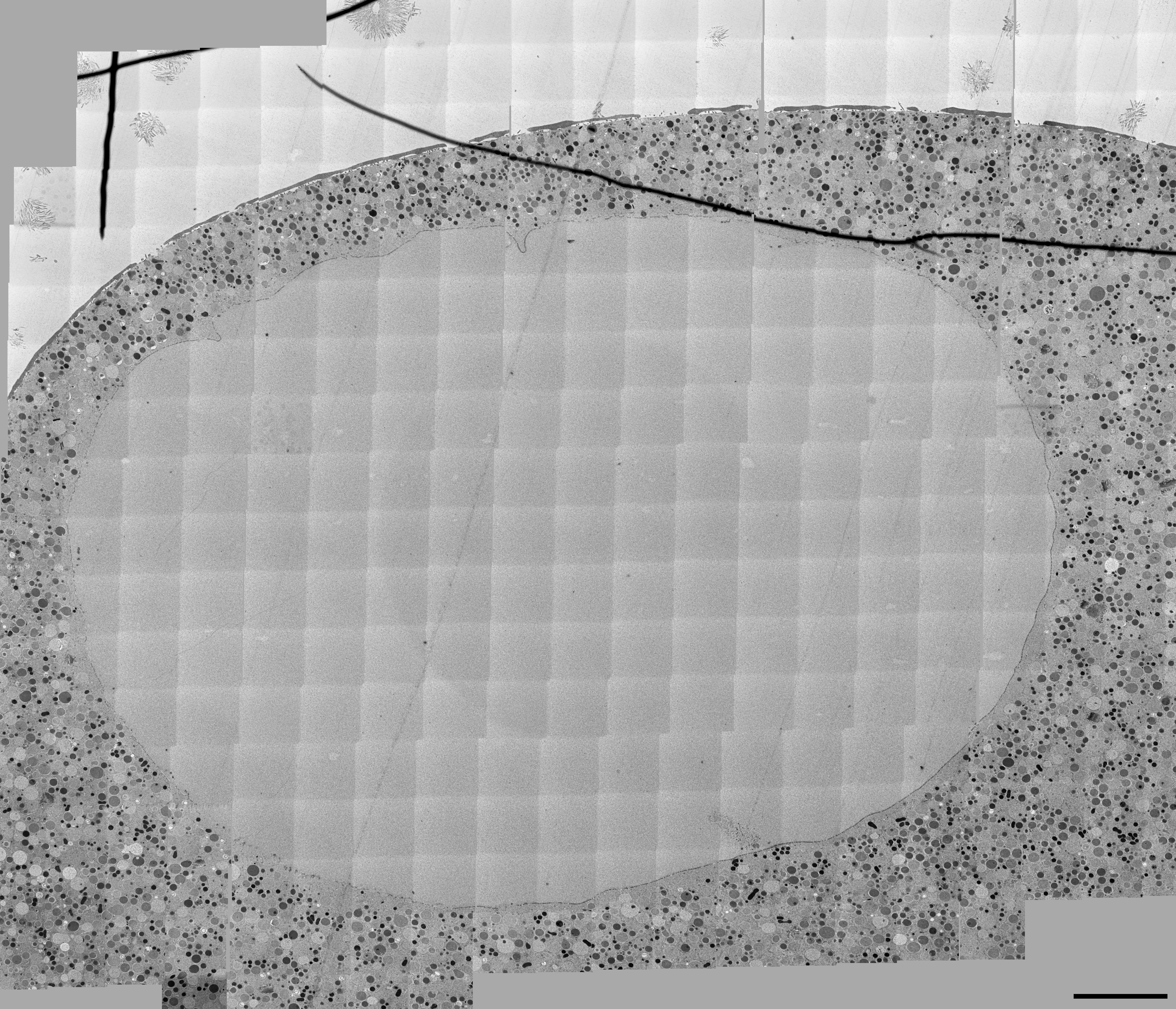

### Supplemental Figure 2

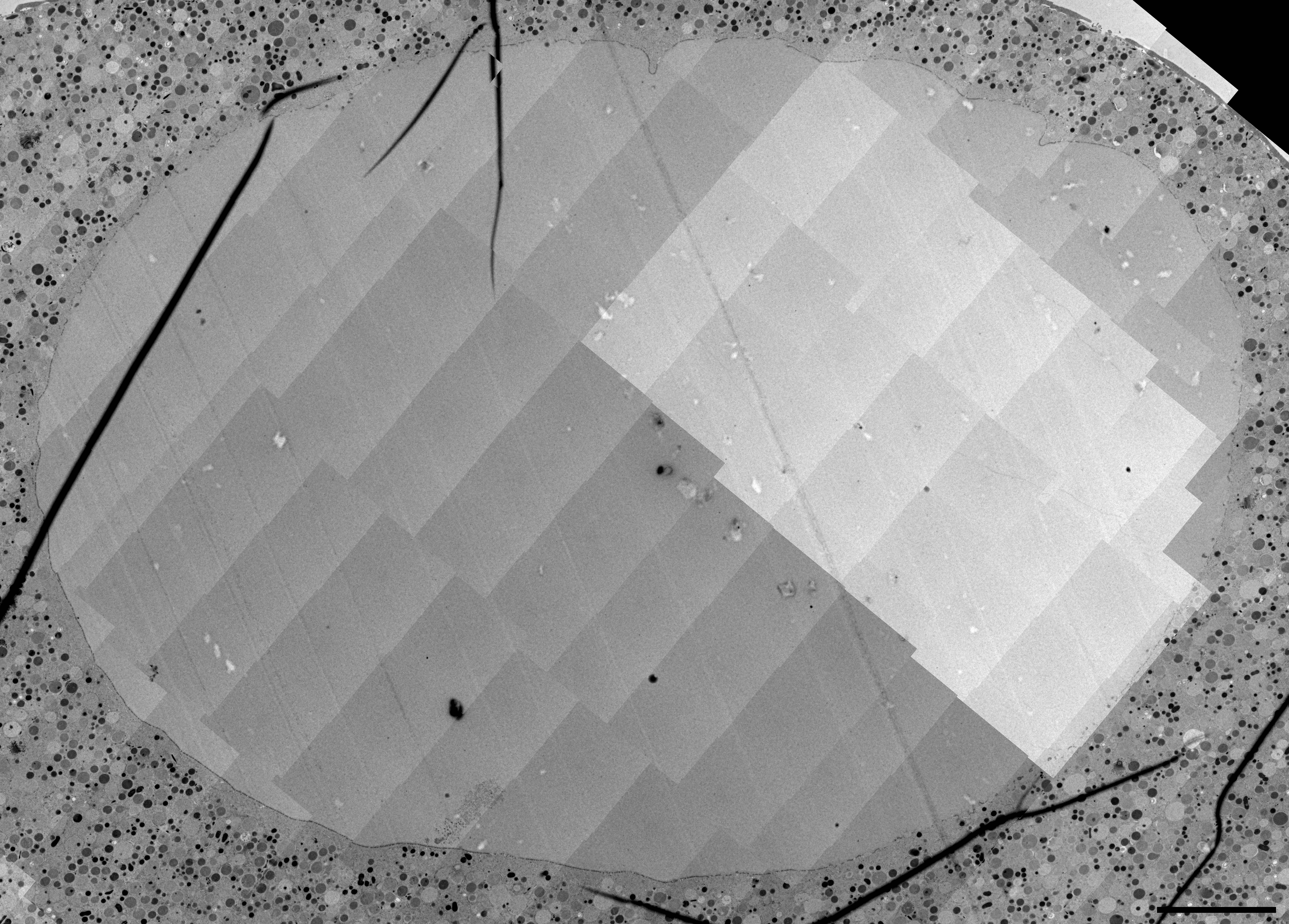

### Supplemental Figure 3

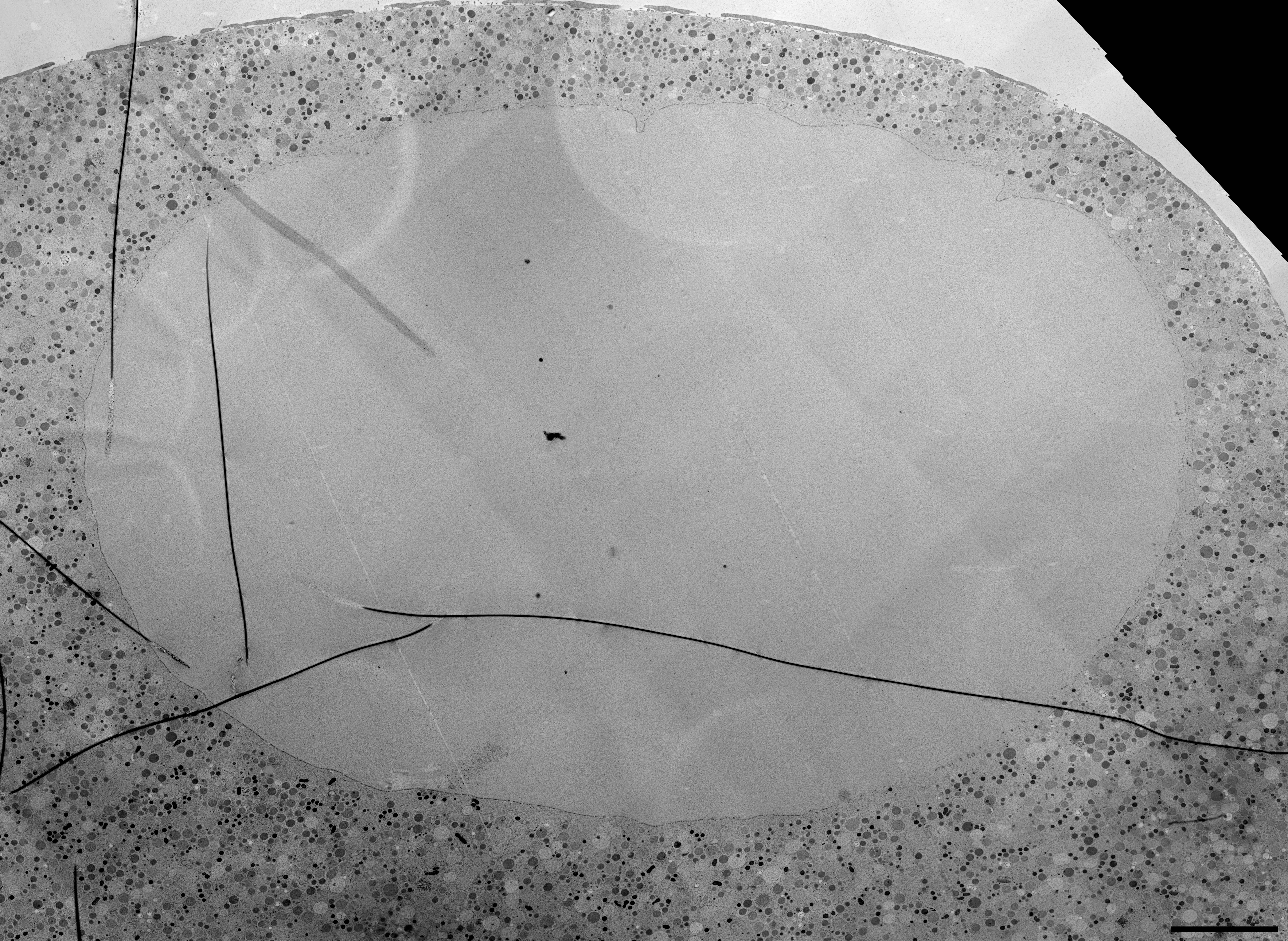

### Supplemental Figure 4

#### Figure S4.

A

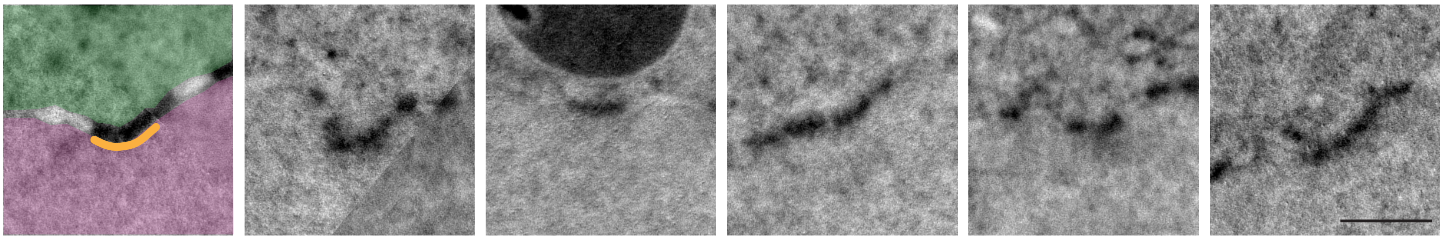

B

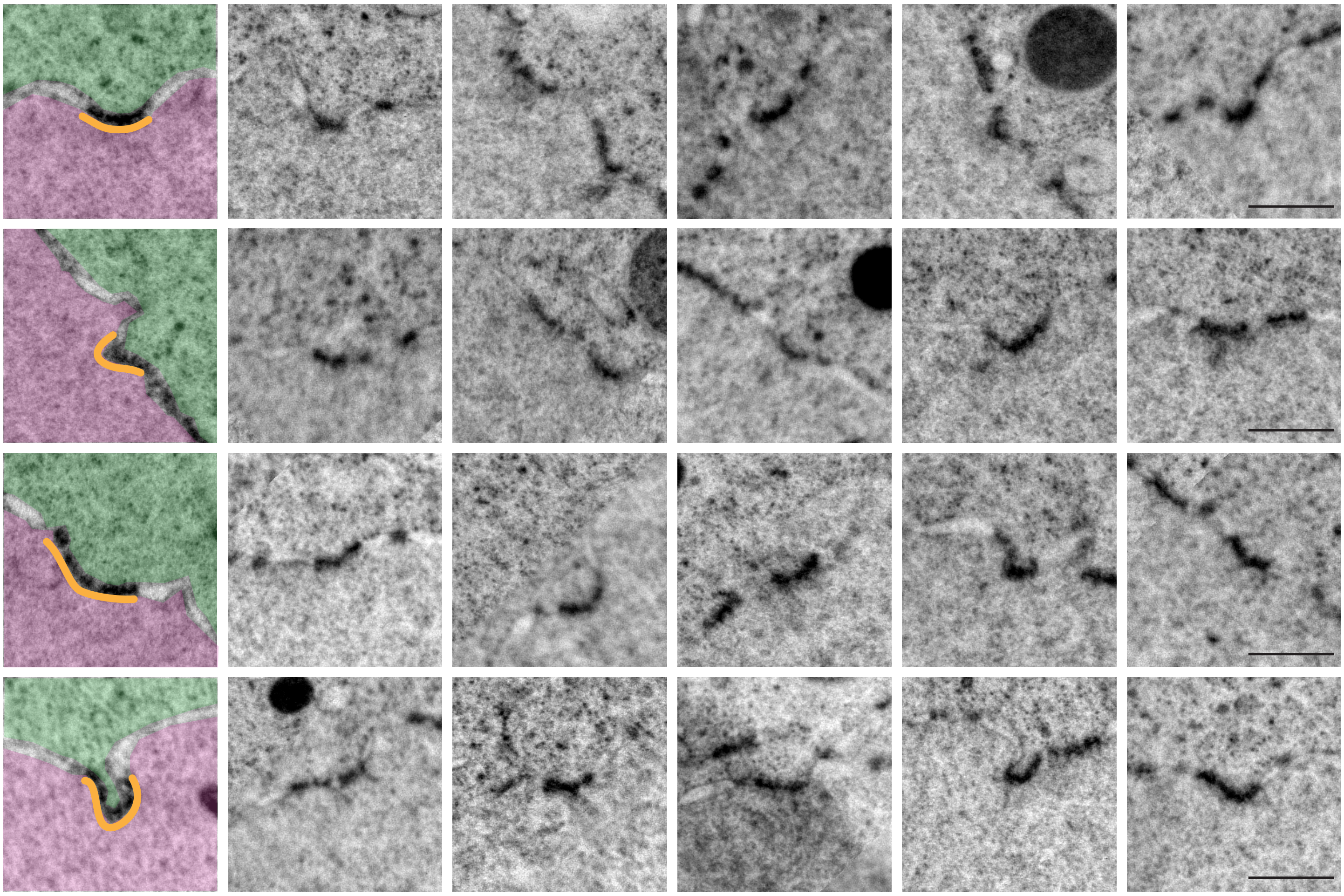

C

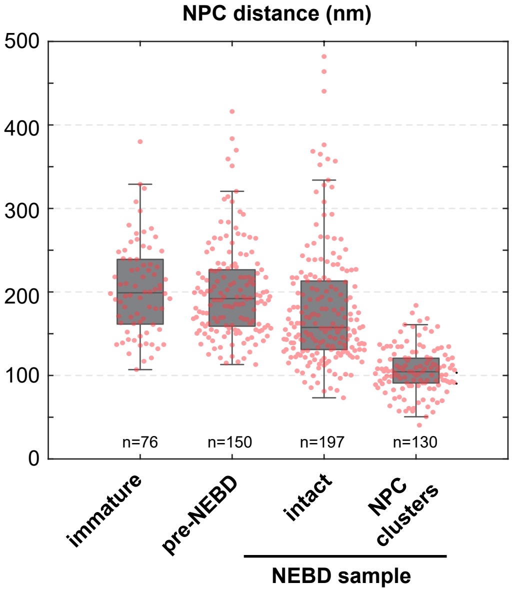
